# The Topological Architecture of Brain Identity

**DOI:** 10.1101/2025.06.20.660792

**Authors:** Simone Poetto, Haily Merritt, Andrea Santoro, Giovanni Rabuffo, Karolina Finc, Demian Battaglia, Francesco Vaccarino, Manish Saggar, Andrea Brovelli, Giovanni Petri

**Affiliations:** Centre for Modern Interdisciplinary Technologies, Nicolaus Copernicus University in Toruń, Poland; CENTAI Institute, Turin, Italy; Program in Cognitive Science, Indiana University, Bloomington, IN, USA; Department of Informatics, Indiana University, Bloomington, IN, USA; Institut de Neurosciences des Systèmes (INS), UMR 1106, INSERM, Aix-Marseille Université, Marseille, France; Laboratoire de Neurosciences Cognitives et Adaptatives (LNCA), Faculté de Psychologie, Université de Strasbourg, Strasbourg, France; Dipartimento di Scienze Matematiche, Politecnico di Torino, Turin, Italy; Department of Psychiatry and Behavioral Sciences, Stanford University School of Medicine, Stanford, CA, USA; Wu Tsai Neurosciences Institute, Stanford University, Stanford, CA, USA; Institut de Neurosciences de la Timone (INT), UMR 7289, CNRS, Aix-Marseille Université, Marseille, France; NP Lab, Network Science Institute, Northeastern University London, London, UK; Department of Physics, Northeastern University, Boston, USA

## Abstract

Accurately identifying individuals from brain activity—functional fingerprinting—is a powerful tool for understanding individual variability and detecting brain disorders. Most current approaches rely on functional connectivity (FC), which measures pairwise correlations between brain regions. However, FC is limited in capturing the higher-order, multiscale structure of brain organization. Here, we propose a novel fingerprinting method based on homological scaffolds, a topological representation derived from persistent homology of resting-state fMRI data. Using data from the Human Connectome Project (*n* = 100), we show that scaffold-based fingerprints achieve near-perfect identification accuracy ( *∼* 100%), outperforming FC-based methods (90%), and remain robust across preprocessing pipelines, atlas choices, and even with drastically shortened scan durations. Unlike FC, in which fingerprinting features localize within networks, scaffolds derive their discriminative power from inter-network connections, revealing the existence of individual mesoscale organizational signatures. Finally, we show that scaffolds bridge redundancy and synergy by balancing redundant information along high-FC border edges with synergistic interactions across the topological voids they enclose. These findings establish topological scaffolds as a powerful tool for capturing individual variability, revealing that unique signatures of brain organization are encoded in the interplay between mesoscale network integration and information dynamics.

---

The human brain is a complex network of hierarchically interconnected regions that continuously exchange information [1–5]. Neuroimaging research has demonstrated that brain connectivity shares common architectural properties across individuals, such as efficient information processing pathways [6], heavy-tailed distributions of connection weights, node degrees with the presence of hubs and rich-club organizations [7–9], modular structures [10], and efficient wiring costs [11]. At the same time, each brain carries unique and subject-specific signatures [12], which are continuously shaped by individual cognitive trajectories, both along healthy agingas well as neurodegeneration [14, 15].

Advances in functional neuroimaging have demonstrated that these individual signatures are highly reliable. Patterns of functional connectivity (FC)—defined as the statistical dependencies between the activity of distinct brain regions [2–5, 16]—are so distinctive they can serve as ‘fingerprints’ for identifying individuals from a group[17, 18].

This functional fingerprinting has since been extended to other modalities, including EEG [19–22], fNIRS [23], and MEG [24, 25], and has been shown to remain robust across different tasks and mental states [26–28]. Clinically, alterations in an individual’s connectivity profile have been proposed as potential biomarkers for tracking progression toward neurodegeneration or cognitive impairment [14, 15, 29, 30].

However the power of FC-based fingerprinting relies on the implicit assumption that brain interactions are primarily pairwise. A growing body of evidence challenges this view. Information-theoretic approaches, such as partial information decomposition [31], have revealed higher-order forms of coordination, including redundancy and synergy among three or more regions [32–35], that pairwise FC cannot capture. Complementary advances in network science and topological data analysis (TDA) have uncovered rich topological structure across modalities, including structural [36], functional [37–39], and representational [40, 41] networks, as well as cortical morphology [42, 43], EEG [44], and PET [45, 46] data. Among TDA tools, the *homological scaffold* [37] is particularly promising: using persistent homology[47], it identifies and aggregates subsets of links forming one-dimensional topological cycles. This approach avoids arbitrary thresholding, preserves essential multiscale structure, and provides a principled topological summary of mesoscale network organization [48, 49]. Scaffold-based analyses have already differentiated global brain state alterations, such as psilocybin versus placebo [37], and revealed shifts in integration patterns across experimental conditions [50] and tasks [35, 51].

Overall, these results strongly suggest that a paradigm shift considering higher-order functional relations and multiscale topological structures is needed for accurate brain fingerprinting. Here, we test the hypothesis that homological scaffolds provide a more accurate representation of individual brain organization with respect to conventional pairwise descriptors. We address this along three directions: (i) evaluating whether scaffold-based representations improve subject identification relative to standard FC; (ii) examining the anatomical distribution of discriminative information, specifically whether the most informative scaffold edges cluster within canonical systems or bridge heterogeneous networks; and (iii) assessing whether scaffold cycles capture regions of elevated synergistic information exchange.

Using test–retest resting-state fMRI data from the Human Connectome Project, we show that scaffolds achieve near-perfect fingerprinting performance with only a fraction of the full connectome. The most discriminative scaffold edges predominantly span across networks rather than within them, suggesting that subject-specific signatures arise from unique patterns of inter-network integration rather than within-network coherence. Finally, we demonstrate that topological cycles consistently enclose regions characterized by high synergy in information flow. Taken together, these results establish homological scaffolds as a compact, robust, and interpretable framework for capturing individual brain identity, opening new perspectives on the interplay between network topology and information dynamics in the human connectome.

## RESULTS

To investigate individual differences in brain functional organization, we analyzed resting-state fMRI data from 100 participants, each scanned in two separate sessions as part of the Human Connectome Project (HCP) [52]. FC matrices were constructed for each participant and session by computing Pearson correlations between the time series of 300 brain regions defined by the Schaefer atlas [53] (Fig. 1A).

**FIG. 1.**
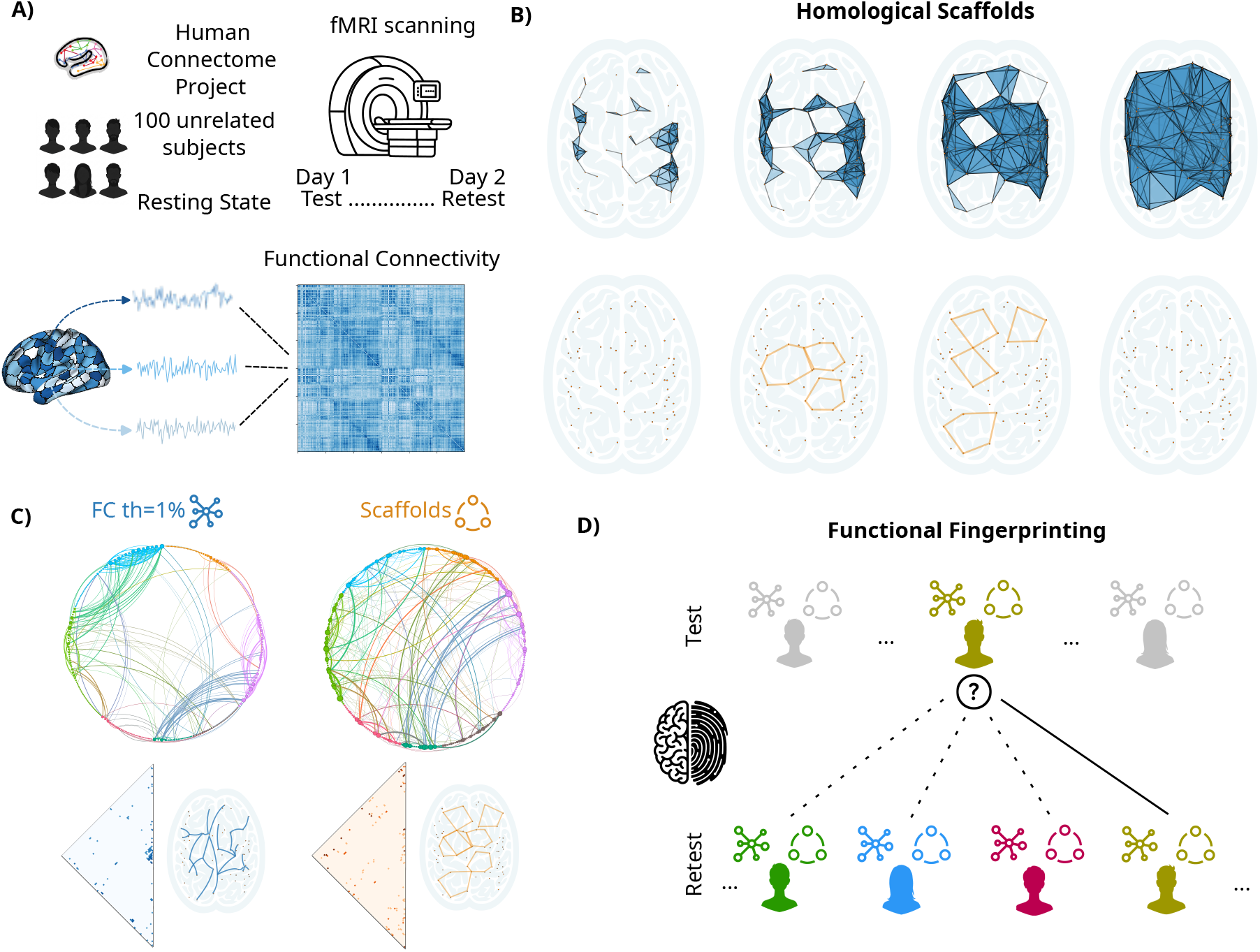
Schematic of scaffold computation and fingerprinting workflow. (A) Resting-state fMRI scans from the 100 unrelated subjects of the Human Connectome Project are used to compute individual functional connectivity matrices from two sessions (*∼* 10 days apart). (B) Conceptual illustration of homological scaffold construction. Edges are progressively added in decreasing weight order, and those participating in topological cycles (*H*_1_ features) form the scaffold. (C) Comparison between a functional connectivity matrix thresholded to retain 1% of the strongest connections (left) and the corresponding scaffold (right), which achieves similar sparsity while preserving topologically informative links. (D) Visual representation of the functional fingerprinting process: an individual’s test session brain connectivity matrix is compared against a database of re-test session matrices to identify the correct match based on the highest similarity.

To test whether topological structure provides more robust subject fingerprints than conventional FC, we applied persistent homology using weighted rank filtration, in which edges are added in decreasing order of their FC weight [54]. During this procedure we tracked the birth and death of topological features, specifically one-dimensional loops (i.e., generators of the *H*_1_ homology group).

We identified the core topological structure by constructing ‘frequency scaffolds’ [37], which consist of all edges participating in at least one homological generator, weighted by the number of different cycles they belong to (Fig. 1B). This procedure typically yields sparse networks, retaining on average *∼* 1% of the original links (Fig. 1C and see Methods for a detailed description). For completeness and as comparative benchmarks, we also computed thresholded FC matrices, retaining the top 25% (following standard FC approaches [55]) and 1% strongest connections (to match scaffold’s densities). Finally, we measured the fingerprinting performance of each representation. We vectorized the upper triangle of the adjacency matrix for each representation, and, for every participant, we computed the Pearson’s correlations between their Session 1 vectors and all Session 2 vectors (and vice versa). Following standard approaches [18, 56], a correct identification occurred when a participant’s own data yielded the highest cross-session correlation. We repeated this procedure in both directions — Session 1 to Session 2 and Session 2 to Session 1 — and defined overall accuracy as the average identification rate across the two directions (Fig. 1D).

### Homological scaffolds outperform functional connectivity in identifying subjects

To assess the performance of the homological scaffold against FC, we first quantified the separation between intra-subject and inter-subject similarity distributions (Fig. 2A). These distributions represent how similar an individual’s brain scans are to their own from a different session (intra-subject) versus how similar they are to the scans of other individuals (inter-subject); a greater separation between these distributions makes it easier to identify a subject uniquely. The scaffold representation demonstrated a markedly clearer separation, with a Cohen’s d effect size (ES = 7.46) that was nearly double that of the FC matrix thresholded at 1% (ES = 3.78) (Fig. 2B). This superior separation translated to a nearly perfect identification rate. While full FC and its various thresholded versions (25% and 1%) plateaued at approximately 90% accuracy, the homological scaffold achieved a 100% success rate for the entire cohort. This perfect identification was consistent in both directions—predicting Session 2 from Session 1 and vice versa (Fig.2A, Section S.1 and Table S1)).

**FIG. 2.**
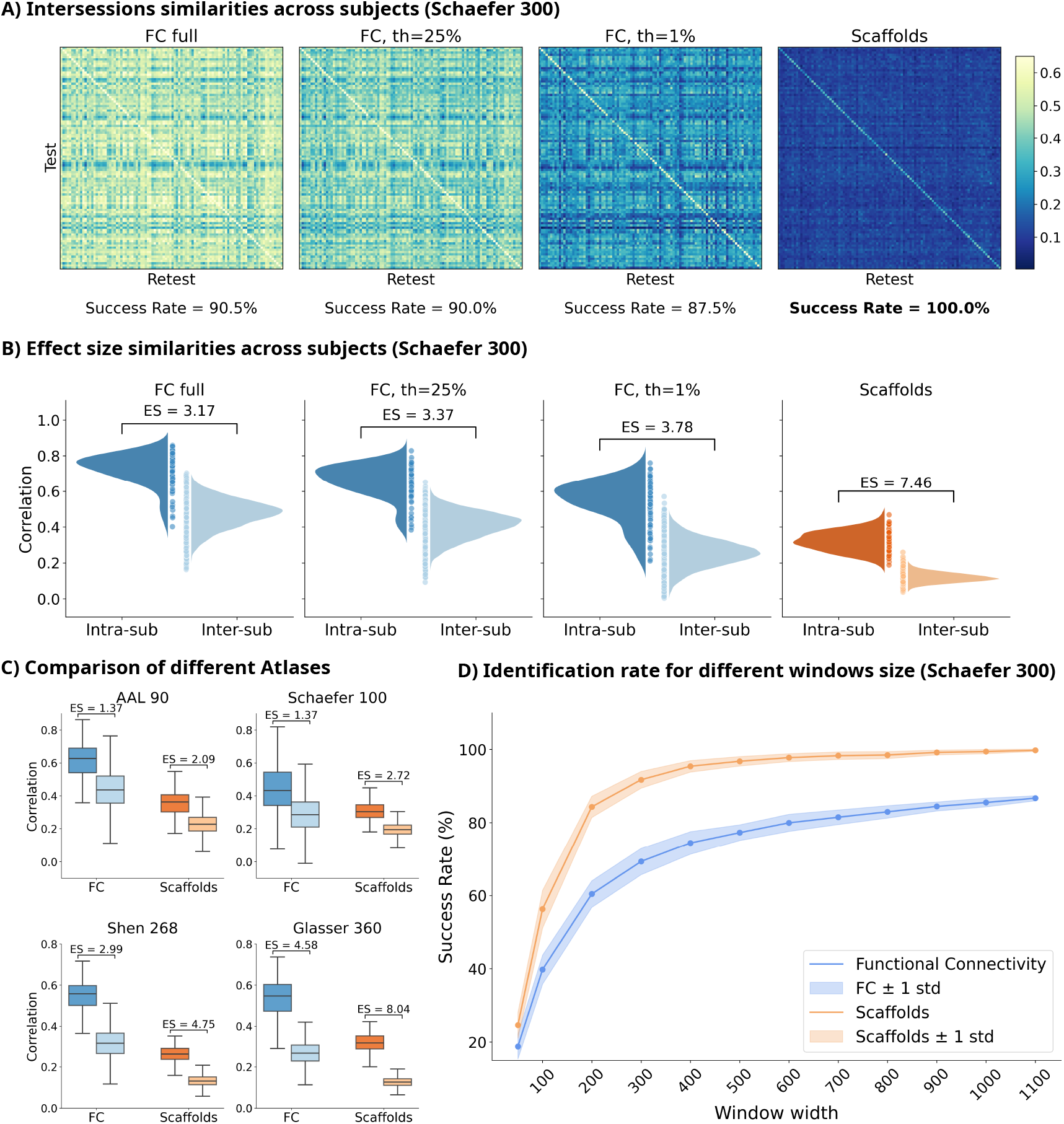
Scaffolds outperform functional connectivity in capturing individual-specific brain patterns. (A) Intersession similarity heatmaps and corresponding identification success rates for full Functional Connectivity (FC) (Success Rate = 90.5%), FC thresholded at 25% (Success Rate = 90.0%), FC thresholded at 1% (Success Rate = 87.5%), and Scaffolds (Success Rate = 100.0%). (B) Violin plots display the distributions of intra-subject (within-subject) and inter-subject (between-subject) Pearson correlations for the four representations shown in (A). Cohen’s *d* values (ES) quantify the separation between these distributions: FC full (ES=3.17), FC th=25% (ES=3.37), FC th=1% (ES=3.78), and Scaffolds (ES=7.46). (C) Boxplots compare intra-subject and inter-subject correlation distributions for FC and Scaffolds across different brain atlases: AAL 90 (FC ES=1.37, Scaffolds ES=2.09), Schaefer 100 (FC ES=2.12, Scaffolds ES=2.72), Shen 268 (FC ES=2.99, Scaffolds ES=4.75), and Glasser 360 (FC ES=4.58, Scaffolds ES=8.04), with their respective Cohen’s d effect sizes indicated. (D) Identification success rate (%) for Functional Connectivity and Scaffolds as a function of fMRI window width (from 100 to 1100 TRs). Shaded areas represent ±1 standard deviation. This demonstrates that scaffold-based representations maintain high discriminability even with limited data, outperforming FC across all window lengths.

Next, we investigated the robustness of these findings. First, we assessed the impact of common fMRI prepro-cessing choices on scaffold performance (Sec. S.2 and Table S4). Overall, we found that fingerprinting capacity of scaffolds is weakly affected by different preprocessing steps. Indeed, the only step to significantly impact performance is low-pass filtering, the omission of which substantially degraded accuracy (from 99% to 83.5% using Glasser atlas data). Conversely, other common preprocessing steps, like global signal regression (GSR), highpass filtering, and the choice of scan acquisition order (left-right vs. right-left phase encoding) have minimal or slightly detrimental effects on scaffold identifiability, which remained at or near 100% in most conditions.

Second, we evaluated fingerprinting performance for different brain atlases, including anatomical (AAL90 [57, 58]) and functional parcellations of varying resolutions (Schaefer100/300 [53], Shen268 [59], Glasser360 [60]) (Fig. 2C, Sec. S.1 S.1, Table S2). Across all atlases, scaffolds consistently outperformed the corresponding 1% thresholded FC matrix (as well as the 25% thresholded and full FC, see SI). Identifiability scores strongly depended on atlas resolution, increasing from 43% (AAL90) and 64.5% (Schaefer100) with scaffolds to 100% (Schaefer300) and 99.5% (Glasser360). As also shown in previous studies [61], this suggests that higher resolution parcellations generally capture better subject-specific details.

Finally, we examined how scaffold performance changed with shorter scan lengths.

Specifically, we considered progressively shorter windows of fMRI data, from 1200 TRs ( *∼* 14.4 minutes) to 50 TRs ( *∼* 0.6 minutes), and computed the identification success rate at each step to quantify identification accuracy (Fig. 2D, see Methods for details, and Fig. S2 for results on effect size).

Across all window lengths, scaffolds again consistently outperformed thresholded FC, maintaining a high effect size comparable to that achieved using the full FC timeseries, even for windows as short as 100 TRs.

This indicates that scaffolds capture stable individual traits even at shorter time scales or limited scan time. In Sec. S.3 we also provided a comparison of fingerprinting capacity within- and across-sessions, showing that within the same session the fingerprinting capacity at short timescales of FC and scaffolds is comparable, supporting the idea that scaffolds captures preserved functional traits, while FC better captures transient states.

### Scaffold edges are distributed across the brain

Having established the superior accuracy and robustness of scaffolds, we next asked: *what features underlie their effectiveness*? In particular, we investigated where individual-specific information is concentrated within scaffold networks. To do this, following Finn et al. [18], we calculated the *differential power* for each edge in both the FC (1% thresholded) and scaffold representations. This metric identifies edges that are highly stable within an individual across sessions but highly variable between individuals. Following previous work [18], we selected the top 0.2% of edges with the highest differential power, considering these as the most critical for fingerprinting.

We then examined the distribution of these high-importance edges across 7 canonical resting-state networks [62]. A marked difference emerged (Fig. 3A-B): for FC, the vast majority (72%) of important edges were within individual networks (intra-network connections). In contrast, for the scaffold, the majority (56%) of important edges spanned different networks (inter-network connections). This suggests that –while FC-based identity is reflected in local functional network properties– scaffold-based identity relies on the integration across networks, reflecting a mesoscale organization.

**FIG. 3.**
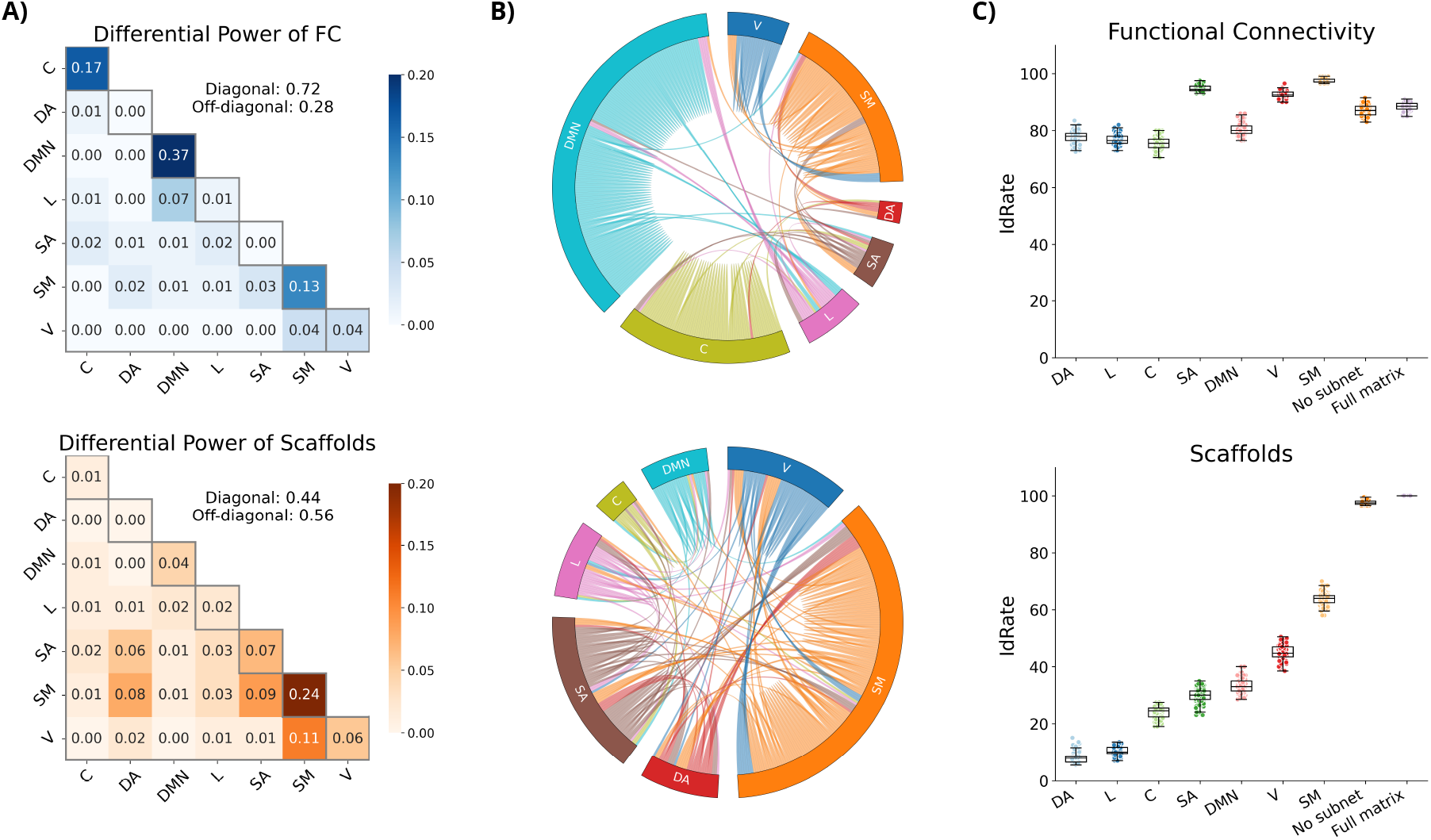
Scaffolds rely on inter-network connectivity to enhance individual identification. (A) Heatmaps illustrating the differential power of edges for FC (top) and scaffolds (bottom). In FC, 72% of differential power is concentrated within intra-network connections (diagonal: 0.72), while 28% spans across inter-networks (off-diagonal: 0.28). In contrast, scaffolds distribute differential power more evenly, with 44% within-network (diagonal: 0.44) and 56% across networks (off-diagonal: 0.56), involving all 7 canonical resting-state networks. (B) Circular connectivity diagrams (connectograms) illustrating the location of high differential power edges in FC (top) and Scaffolds (bottom). (C) Box plots comparing identification rates (IdRate) for FCs and Scaffolds when considering only edges across different subnetworks ‘No subnet’) versus using only intra-network edges from specific subnetworks (DA: Dorsal Attention, C: Control, DMN: Default Mode Network, L: Limbic, SA: Salience/Ventral Attention, SM: Somatomotor, V: Visual) or the full matrix. This further confirms that FC heavily relies on within-network connections, while scaffold identity relies on inter-network connections.

To directly test this hypothesis, we repeated the finger-printing analysis using only intra-network edges or only inter-network edges for FC and scaffolds (Fig. 3C, Section S.1 S.2, Table S3). For FC, intra-network edges alone supported relatively high identification rates (e.g., 73% using Default Mode Network edges), often comparable to full-network performance. This reinforces the view that FC-based identity signatures are localized within specific brain systems. In contrast, scaffold-based identification suffered when restricted to intra-network edges, with performance dropping to 10–50% across networks (maximally 60% for the Somatomotor network). Strikingly, using only the inter-network scaffold edges restored performance to nearly full accuracy ( *∼* 98%), highlighting again that the key individual-specific information in scaffolds lies in the links between networks. Scaffolds thus appear to capture an inherently mesoscale signature of individual brain organization.

### Scaffold edges balance synergy and redundancy

Having established the scaffold’s discriminative power, we next investigated its structural properties as a higher-order object. So far, we have considered its individual links, but the scaffold’s fundamental building block—an *H*_1_ generator—is a closed loop of highly connected edges. This structure defines two distinct types of connections (Fig 4B). The border links are the high-correlation edges that form the cycle’s perimeter; by construction of the filtration, they are among the strongest links in the network. In contrast, *H*_1_ generators are also characterized by a set of internal links that span the void enclosed by the border. These links are ‘missing’ when the cycle forms because their correlation is lower than that of the border edges.

**FIG. 4.**
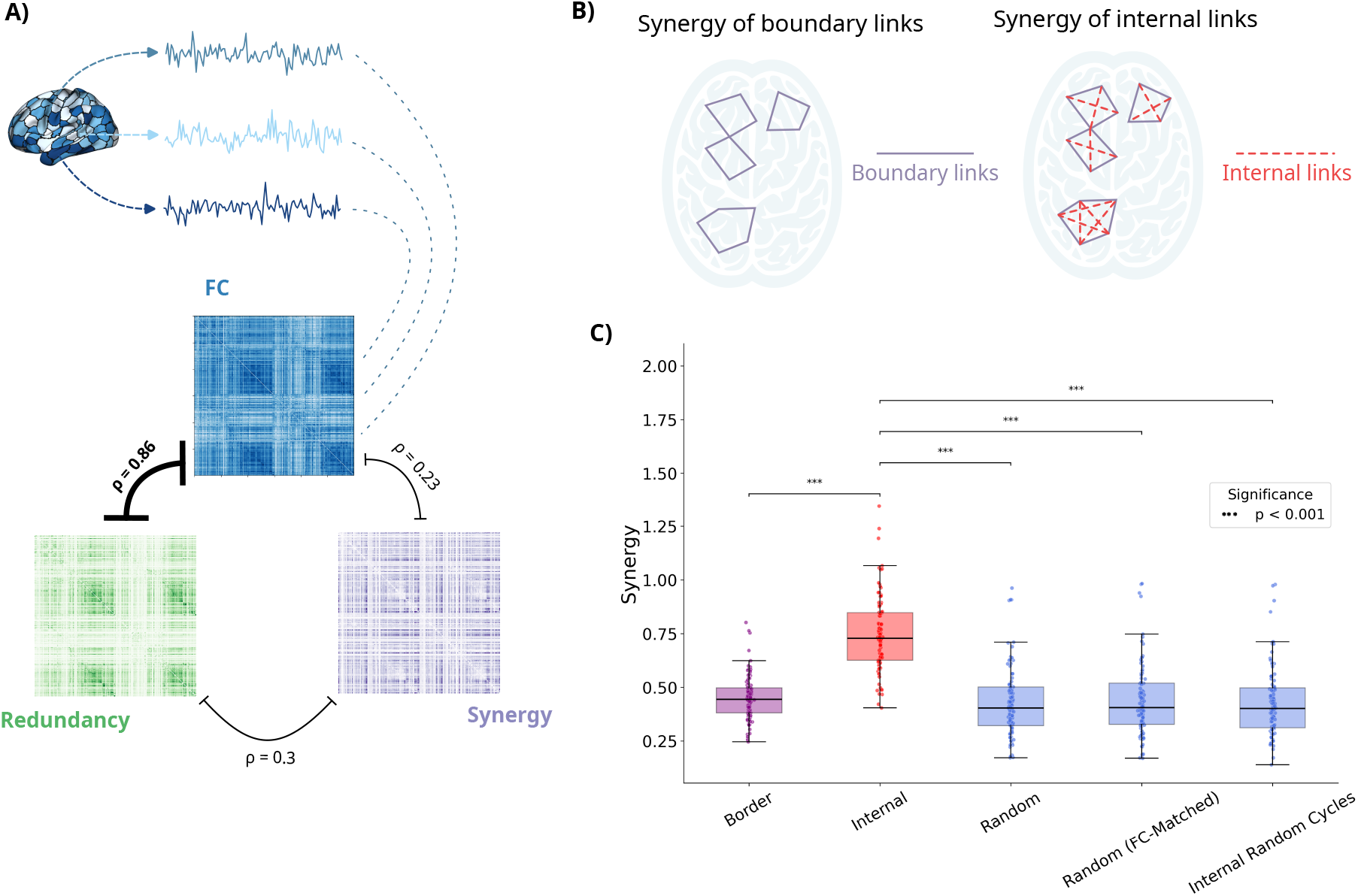
Information-theoretic properties of homological cycles. (A) Diagram illustrating the relationships between FC, redundancy, and synergy across brain regions. FC is strongly positively correlated with redundancy (*ρ*=0.86), but only weakly with synergy (*ρ*=0.23). Redundancy and synergy also show a weak correlation (*ρ*=0.3). (B) Conceptual illustration distinguishing *border* links — edges that form the borders of topological cycles (solid lines) — from *internal* links that span the interior of the topological cycles (dashed red lines). (C) Boxplots comparing mean synergy (in bits) across different categories of links: Border (true scaffold border links), Internal (true internal links spanning topological holes), Random (randomly selected edges), Random (FC Match) (randomly selected edges matched to FC correlation values of internal links), and Random Cycle Internal (internal edges from randomly selected cycles). Internal links show significantly higher synergy than border links and random selections, suggesting a unique role in integrative information processing within topological cycles.

Motivated by this distinction, we investigated the information-theoretic properties of these two edge types. Following prior works [31], we adopted the integrated information decomposition (Φ-ID) framework [63] to compute subject-specific synergy and redundancy matrices for every pair of brain regions. We first confirmed that functional connectivity (FC) is strongly correlated with redundancy, while synergy shows no significant relationship with FC (Fig. 4A). This implies that the high-FC border links primarily capture redundant information. A natural question, therefore, is how synergy behaves with respect to these topological cycles. We hypothesized that the internal edges, despite their lower correlation and redundancy, support greater synergistic interactions, acting as integrative ‘bridges’ across the topological holes they span.

To test this, we compared the synergy values of internal edges versus border edges. Indeed, we found that internal links displayed significantly higher synergy than border links (Fig. 4C). To confirm this result is not a byproduct of low correlation alone (i.e., an inverse relationship between synergy and redundancy), we compared the synergy of internal edges against three null distributions: (i) one obtained by sampling synergy values randomly from all edges, (ii) a second one obtained by sampling synergy values only from edges with FC/redundancy values matched to those of the internal links, and (iii) a third one obtained by sampling synergy values from random non-topological closed cycles (to assess whether the effects emerged from pure geometrical effects [64]). In all cases, the internal scaffold edges showed significantly higher synergy than expected by chance (Fig. 4D).

Together, these findings uncover a novel link between the topological organization of brain networks and their information-processing profiles. Scaffold representations delineate a functional architecture in which redundant interactions concentrates along the borders of topological cycles, while synergistic interactions preferentially occupy the voids they enclose. This spatial segregation of redundancy and synergy suggests that scaffolds encode a balance between structured, stable pathways and flexible, integrative processing. Such a balance may underlie the emergence of individual-specific scaffold signatures, capturing distinctive aspects of brain organization that remain hidden to conventional FC analyses.

## DISCUSSION

Identifying individuals based on their unique patterns of brain activity and connectivity — brain fingerprinting — has emerged as a powerful paradigm for understanding individual differences in brain organization [17, 18]. While functional connectivity (FC) derived from fMRI has proven remarkably effective for this purpose [18, 65], it primarily captures pairwise linear correlations, potentially overlooking richer, higher-order organizational principles. Here, we introduced an alternative approach leveraging topological data analysis (TDA), specifically homological scaffolds [37], to extract a unique fingerprint based on the persistent topological structure of the functional connectome. Our findings demonstrate that these topological fingerprints not only achieve near-perfect identification accuracy, but also exhibit superior robustness and reveal distinct organizational features compared to traditional FC-based methods.

The superiority of these topological measures distilled from conventional correlation-based measures aligns with emerging evidence that brain organization is fundamentally shaped by wiring constraints and communication efficiency principles [11, 66]. In this perspective, topological cycles may represent optimal routing paths that minimize metabolic costs while max-imizing information integration—constraints that could vary systematically across individuals due to genetic and developmental factors [67], as hinted also by the recent finding that scaffold-derived task flexibility is modulated by dopamine-transporter (DAT) availability, with lower DAT density supporting greater reconfiguration across tasks [35]. This suggests that individual differences in topological fingerprints may, at least in part, reflect underlying neuromodulatory architectures that shape the balance between stability and flexibility in large-scale brain dynamics.

### Topological fingerprinting

The central result of our study is the striking superiority of homological scaffolds over standard FC measures for individual identification. Using test-retest resting-state fMRI data from the Human Connectome Project, scaffolds achieved 100% accuracy in identifying 100 unrelated individuals, significantly outperforming full FC and thresholded FC matrices, which plateaued around 90% (Table S1, Fig. 2A). This advantage was not incidental; it proved remarkably robust across various standard preprocessing choices (Table S4), multiple brain atlases of differing resolutions and modalities (Table S2, Fig. 2C), and perhaps most notably, across substantially shortened time series lengths (Fig. 2D). The scaffolds’ ability to maintain high discriminability even with data segments as short as 100 TRs ( *∼* 72 seconds) suggests they capture highly stable, intrinsic topological traits of an individual’s functional brain architecture, that are not explained by the modest correlations with structural and functional connectomes [51], and thus potentially offering advantages for clinical applications or studies with limited scan time. This is consistent with recent studies on the temporality of brain fingerprinting, which demonstrated that bursts of ‘identifiability’ can occur even over short time scales [68, 69] (and the additional results reported in Sec. S.3).

### Scaffolds as a distributed individual functional watermark

But, why are scaffolds such effective fingerprints? Our results suggest that scaffolds capture fundamentally different aspects of collective brain dynamics compared to FC. When examining the edges most critical for identification using differential power (DP) [18], we found a clear distinction: high-DP FC edges were predominantly located within canonical brain networks (72%), particularly the default mode, control, and visual systems, consistent with previous reports emphasizing the role of fronto-parietal and default mode networks in FC-based identifiability [18, 70]. In stark contrast, high-DP scaffold edges were predominantly found between different networks (56%) (Fig. 3a,b). This finding echoes recent work on higher order interactions in brain networks that emphasizes integration across canonical functional systems [71].

Furthermore, restricting the fingerprinting analysis to only intra-network edges severely degraded scaffold performance, whereas using only inter-network scaffold edges largely recovered the full scaffold’s near-perfect accuracy (Fig. 3c). This indicates that the individual identifying power of scaffolds resides in their ability to capture the mesoscale distributed organization of brain function – the specific pattern of topological loops and holes formed by interactions across large-scale systems, rather than the localized connectivity strength within those systems. This finding resonates with recent work suggesting that individual differences in cognition are better predicted by between-network rather than within-network connectivity [72, 73]. The inter-network edges captured by scaffolds may reflect individual variations in network integration capacity—a key feature of cognitive flexibility and executive function [74, 75]. This mesoscale topological signature appears to be a more unique and stable individual identifier than the strength of local connections typically emphasized by FC.

### A bridge between topology and information theory

Homological scaffolds emerge as a first explicit link between topology and multivariate information theoretic description. In line with previous work [31], we found FC strength strongly correlated with redundant information between regional time series (Fig. 4A).

The scaffold construction selects for edges involved in forming topological cycles (*H*_1_ generators). By definition of the weighted rank filtration, edges forming the ‘borders’ of these cycles have high FC weights and, consequently, high redundancy. The intriguing finding emerged when examining the *internal* edges that span the topological voids enclosed by these borders – edges characterized by relatively lower FC and redundancy. These internal edges exhibited significantly higher synergistic information than the border edges, and significantly higher than expected by chance, even when controlling for FC/redundancy levels (Fig. 4C). This sug-gests a novel link between network topology and information dynamics: the 1-dimensional ‘holes’ revealed by persistent homology are not merely absences of strong pairwise correlation, but are structured regions characterized by heightened synergistic interactions among the connections spanning them.

This interplay between redundancy and synergy within the scaffold structure offers a potential explanation for its discriminative power. The border links demonstrate high redundancy, with strong pairwise correlations indicating shared informational content. In contrast, considering pairs of links that span the internal loops reveals high synergy between them, an interaction that emerges despite the weak correlation characterizing the nodes. This spatial segregation of information modes parallels recent findings that synergistic interactions support conscious awareness and cognitive flexibility [31, 71]. The topological cycles may thus delineate functional units where redundant ‘backbone’ connections provide stable information channels, while synergistic ‘bridges’ enable flexible recombination of information—a balance that could be highly individual-specific due to its role in supporting each person’s unique cognitive style [76].

An important practical advantage of scaffolds is their typically high sparsity (*≈* 1% of total edges), which makes them computationally efficient for large-scale studies. This efficiency, combined with their robustness to short scan durations, positions scaffolds as a practical tool for clinical implementation where scan time is limited. While strategic study design [77], broader sampling [78], and scan duration [79] are often thought to improve predictive power, these results, in conjunction with others [80], suggest novel parsimonious and data-efficient routes for identifying individual differences in MRI signal.

Importantly, when we repeated the analysis *within* each session (Fig. S3-S4), the gap between scaffolds and FC decreased notably: FC success rates and effect sizes rose to scaffold-like levels, whereas scaffold performance was unchanged. This convergence indicates that FC fingerprints derive a substantial boost from transient, state-dependent co-activation patterns that are naturally aligned within the same scan. By contrast, the scaffold signal—anchored in persistent topological cycles—appears largely insensitive to such momentary fluctuations, capturing instead trait-like organizational features that generalize across days.

### Limitations and future work

Our study naturally presents some limitations. First, the scaffolds were derived from static, time-averaged FC matrices. While the sliding window analysis showed robustness over shorter intervals, exploring scaffolds derived directly from dynamic connectivity or using time-resolved topological methods could reveal further insights into transient brain states and their individual specificity. Second, the interpretation of information-theoretic measures like synergy and redundancy in the context of BOLD fMRI is still an active area of research [31, 81]. While our findings point to a compelling link between topology and synergy, the precise functional meaning requires further investigation. Third, our analysis focused on *H*_1_ homology (loops); exploring higher-dimensional topological features (*H*_2_, cavities) might uncover additional organizational complexity [36, 40, 64, 71].

Future research should investigate the functional relevance of these topological fingerprints. For example, relevant questions are: How do individual variations in scaffold structure relate to cognitive abilities, personality traits, or behaviour? Can changes in scaffold topology track learning, development, or ageing? Additionally, combining scaffolds with task-based fMRI could reveal how individual topological signatures reconfigure during cognitive demands, potentially linking stable traits to state-dependent flexibility [82, 83]. Furthermore, applying topological fingerprinting to clinical populations could be particularly fruitful. Alterations in brain connectivity are hallmarks of numerous neurological and psychiatric disorders [84, 85]. Recent work has shown altered topological properties in autism [86], schizophrenia [87], and Alzheimer’s disease [88]. Our scaffold approach might be particularly sensitive to these alterations given its focus on mesoscale organization, which appears disrupted across multiple psychiatric conditions [89]. Investigating whether scaffold properties are differentially affected in these conditions could provide novel biomarkers for diagnosis, prognosis, or treatment response, potentially offering greater sensitivity than traditional FC measures due to their robustness and mesoscale nature. In conclusion, this work introduces homological scaffolds as a potent and robust method for brain fingerprinting, significantly surpassing traditional functional connectivity approaches. By capturing the persistent topological structure of brain interactions, particularly the mesoscale arrangement of connections spanning across large-scale networks, scaffolds provide a unique window into individual brain organization. Our findings linking the topological voids within scaffolds to heightened synergistic information suggest a deeper connection between network topology and information processing principles. This work highlights the power of moving beyond pairwise correlations [90] by joining tools from topology and information theory [51, 91] to unravel the complex, individualized architecture of the human brain.

## I. METHODS

### A. Dataset

We utilized data from 100 unrelated healthy young adults (54 females, 46 males, mean age = 29.1*±*3.7 years) provided in the Human Connectome Project (HCP) 900-subject data release [52, 92]. The HCP consortium curated this specific subset to ensure individuals were not family relatives, which was crucial for our study to avoid potential confounds related to family structure in identifiability analyses. All participants provided written informed consent according to the HCP protocol, which was approved by the local Institutional Review Board at Washington University in St. Louis. All experiments were performed in accordance with relevant guidelines and regulations.

We focused on the resting-state fMRI data (HCP filenames: rfMRI REST1 and rfMRI REST2). These were acquired in separate sessions on two different days. Each session included scans with both left-to-right (LR) and right-to-left (RL) phase-encoding directions to mitigate susceptibility distortions. For all primary analyses, we exclusively utilized the data from the LR phase-encoding direction. To validate this approach, we conducted a verification analysis where we compared the functional connectivity (FC) matrices from the LR scans alone against the mean FC matrices obtained by averaging the results from the LR and RL scans. The results were highly comparable, confirming that using only the LR data provided a reliable measure of functional connectivity representative of the full dataset. Full details on the HCP restingstate acquisition can be found in [93].

### B. Preprocessing

The data were processed using the HCP minimal preprocessing pipelines [94]. The preprocessing workflow corrected for gradient distortion, head motion, and B0 field inhomogeneities. Functional data were registered to the individual’s T1w structural image and then transformed into MNI152 standard space. All transformations were concatenated and applied in a single step to minimize interpolation blurring, with the final data resampled to 2mm isotropic voxels. To preserve fine-grained spatial detail, no spatial smoothing was applied. Structured noise was removed using ICA-FIX [95, 96], which identifies and removes non-neural signal components. The resulting cleaned time series from the *MNInonlinear* folder served as the primary input for all subsequent analyses.

### C. Parcellation

To define network nodes, we applied several standard brain atlases to the preprocessed fMRI data in MNI space. These included: the Automated Anatomical Labeling atlas (AAL, 90 regions) [57], the Schaefer functional parcellations (using resolutions of 100 and 300 regions) [53], the Shen functional parcellation (268 regions) [59], and the Glasser multimodal parcellation (360 cortical regions, to which we added 19 subcortical areas from the HCP release, for a total of 379 regions) [60, 94]. For each participant and each parcellation, regional time series were extracted by averaging the BOLD signal across all voxels within each defined brain region. For analyses involving brain subnetworks, we utilized the 7 canonical resting-state networks defined by Yeo et al. (2011) [62], assigning each cortical parcel from the Schaefer300 atlas to one of these networks.

### D. Functional Connectivity (FC)

The conventional functional connectivity, denoted as *FC*_*ij*_, was determined by calculating the Pearson correlation coefficient between each pairs *i* and *j* of preprocessed and denoised BOLD signal time series corresponding to a different brain region.

### Pearson correlation coefficient

The correlation coefficient *r*_*ij*_ between two time series *i* = {*i*_1_, …, *i*_*N*_} and *j* = {*j*_1_, …, *j*_*N*_} is defined as:

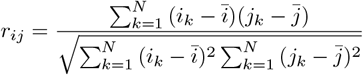

where 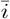 and 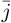 are the means of the time series *i* and *j*, respectively, and *N* is the number of time points. This calculation results in a symmetric *M × M* FC matrix for each subject and session, where *M* is the number of brain regions in the chosen parcellation.

### E. FC Thresholding

To create sparse representations of the functional connectome for comparison, we applied a thresholding procedure to the full FC matrices. This method involves retaining a specific percentage of the strongest connections, ranked by their Pearson correlation values, while setting all other connections to zero. In this study, we used two different thresholds. First, we retained the top **25%** of connections, a common practice [55] in functional connectivity studies. Second, to create a benchmark with a density comparable to that of the homological scaffolds (approximately 1%), we also generated matrices by retaining only the top **1%** of the strongest connections.

### F. Homological Scaffold

To analyze the topological structure of the weighted networks, we used persistent homology [47]. Specifically, we employed the methodology described in [54].

In this framework, a weighted network is converted into a sequence of unweighted graphs, known as a *filtration*. This is achieved by considering the edge weights as the filtration parameter. We build a sequence of simplicial complexes [54, 97], specifically clique complexes, by ordering the network links by their weights in descending order. The filtration is constructed by creating a series of thresholded graphs, where each step includes all edges with a weight greater than or equal to a given threshold. As the threshold is lowered, more edges are included, and the corresponding clique complex grows.

Within this filtration, we track the birth and death of one-dimensional topological holes, which are the generators of the first homology group (*H*_1_). A cycle is *born* at a specific filtration value (i.e., edge weight) when a set of edges forms a closed loop. A cycle is said to *die* when it is filled in by 2-simplices (triangles), which occurs when edges are added that connect the vertices of the cycle, effectively turning the hole into a set of complete subgraphs (cliques). The birth (*β*_*g*_) and death (*δ*_*g*_) of each *H*_1_ generator, *g*, are recorded as indices in the filtration sequence.

To summarize the topological information and understand the importance of individual links in forming cyclical structures, we constructed a *frequency homological scaffold*, as introduced in Petri et al. (2014) [37]. The frequency scaffold is a weighted graph where the edge set is composed of all edges that are part of at least one *H*_1_ generator.

The weight of an edge *e* in the frequency scaffold,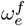, is defined as the total number of distinct *H*_1_ generators to which that edge belongs. This is calculated using the following formula [37]:

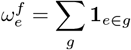

where 1_*e∈g*_ is an indicator function that is equal to 1 if edge *e* is a part of the generator *g*, and 0 otherwise.

All persistent homology calculations were performed using the Ripserer.jl library [98] in the Julia programming language. For each *H*_1_ generator, we choose the representative cycles provided by Ripserer.jl library [98]. While multiple choices are possible, as each homological generator corresponds to a class of equivalent cycles, this choice has been proved to be stable and replicable [37, 49].

### G. Fingerprinting

To assess the ability of different network representations (full FC, thresholded FC, Scaffold) to identify individuals, we implemented a fingerprinting procedure based on cross-session similarity [18]. For each subject and session, the upper triangle of the corresponding connectivity matrix was vectorized.

Let *v*_*i,s*1_ be the vector for subject *i* in session 1 and *v*_*j,s*2_ be the vector for subject *j* in session 2. We computed a similarity matrix **Sim**, where **Sim**_*ij*_ is the Pearson correlation coefficient between *v*_*i,s*1_ and *v*_*j,s*2_. To identify subject *i* from session 1, we found the subject *j*^*∗*^ in session 2 that yielded the maximum similarity: *j*^*∗*^ = arg max_*j*_(**Sim**_*ij*_). Subject *i* was considered correctly identified if *j*^*∗*^ = *i*. This process was repeated to identify subjects from session 2 based on session 1 similarity. The overall *success rate* was calculated as the average percentage of correctly identified subjects across both identification directions (Session 1 *→* Session 2 and

Session 2 *→* Session 1). We provide the disaggregated values in Table S1.

### H. Effect Size

To quantify the discriminability offered by different representations, we calculated the effect size [99], separating the distributions of within-subject and between-subject similarity scores. Within-subject similarity refers to the correlation between vectors of the same subject across the two sessions (i.e., the diagonal elements of **Sim, Sim**_*ii*_). Between-subject similarity refers to the correlations between vectors of different subjects across sessions (i.e., the off-diagonal elements **Sim**_*ij*_ where *i* ≠ *j*). We used Cohen’s *d* as the measure of effect size [100], defined as:

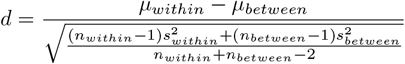

where *µ* and *s*^2^ are the mean and variance of the similarity scores for the within-subject and between-subject distributions, respectively, and *n* is the number of scores in each distribution. The denominator represents the pooled standard deviation.

### I. Bootstrap Subsampling

To assess the stability and robustness of our findings regarding the fingerprinting success rates for FC and scaffolds, we employed a bootstrap subsampling procedure. We randomly sampled 80 subjects without replacement. We then calculated the fingerprinting success rate based only on this subset of 80 subjects. This subsampling process was repeated 1000 times, generating a distribution of success rates, which allowed us to estimate the variability of the metric due to subject selection.

### J. Sliding Window Analysis

To investigate the temporal dynamics and stability of fingerprinting features, analyses were conducted on various shorter portions of the original 1200 TRs restingstate time series. We explored time series of the following lengths: 1100, 1000, 900, 800, 700, 600, 500, 400, 300, 200, 100, and 50 TRs. For each selected time series length, we employed a sliding window approach. Each time series was partitioned into consecutive, potentially overlapping segments of a fixed length, *w*. Each window was treated as a separate realization from which an independent FC matrix was derived. For each length we selected a window spanning from the first TRs *t* = 1 to *t* = *w*. The window was then shifted forward by a step size *s*. A variable sliding parameter *s* was used, with a minimum value of 100 TRs. This meant that for longer time series, there was an unavoidable overlap, resulting in a maximum number of windows. For example, with a time series length of 1100 TRs, only two windows of that length could be selected: one from 0 to 1100 TRs, and a second from 100 to 1200 TRs. When possible, we aimed to select six windows of each length, minimizing the overlap between them or selecting the most spread-apart windows to ensure broad coverage of the original time series. This iterative process continued until the window encompassed the final portion of the time course, resulting in a collection of connectivity matrices. To compare different window lengths, we created numerous realizations. For each realization, we randomly selected a single window per subject and calculated relevant metrics, such as the effect size and success rate, from the derived FC matrices. We repeated this random selection and metric computation process 100 times to ensure the statistical robustness of our findings.

### K. Differential Power

To identify edges most critical for individual identification, we calculated the differential power (DP) for each edge, inspired by the approach of [18]. The differential power metric quantifies the contribution of an edge to fingerprinting by assessing its stability within an individual across sessions relative to its variability across the group. Conceptually, edges with high differential power exhibit connection strengths that are highly consistent for the same person across time but differ substantially between individuals. We used this metric to rank edges and select the top fraction (in this case 0.2%) deemed most important for identification for subsequent analyses, such as computing their densities across canonical brain networks.

### L. Integrated Information Decomposition

To explore the information-theoretic properties of functional connections, we utilized the framework of Partial Information Decomposition (PID). Specifically, we employed the Integrated Information Decomposition (Φ-ID) formalism [63], an extension designed to analyze multivariate time series. This approach decomposes the Time-Delayed Mutual Information (TDMI)—the information shared between the past and present states of two regional time series—into distinct, non-overlapping components: redundancy (*I*_*red*_, information available from both past variables), synergy (*I*_*syn*_, information accessible only from the combination of past variables), and unique information (*I*_*unq,X*_, *I*_*unq,Y*_ ).

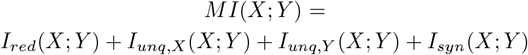

Following prior applications of PID to neuroscientific data [31, 35], we computed pairwise synergy and redundancy for the BOLD time series of every pair of brain regions. For continuous variables, which serve as an adequate model for fMRI time series, this decomposition is achieved by defining redundancy using the Minimum Mutual Information (MMI-PID) principle. This procedure yielded subject-specific synergy and redundancy matrices for each participant. All computations were performed on HRF-deconvolved BOLD signals using the Gaussian copula estimator, as implemented in the hoi software library [101].

### M. Synergy Distribution Analysis

We specifically investigated the synergy values associated with the topological structures identified by the scaffold. We compared the distribution of synergy values for edges belonging to the scaffold *borders* (i.e., edges that are part of the frequency scaffold matrix *S* with *S*_*ij*_ *>* 0) against the synergy values of *internal* edges. Internal edges were conceptually defined as those edges connecting pairs of nodes that lie within a topological hole (an *H*_1_ cycle) but are not part of the cycle boundary itself. To verify that the observed higher synergy of the internal scaffold was not due to confounding factors, we compared its distribution to that of randomly selected edges defined in several ways. All comparisons were within each subject.

1. *Random*: Synergy values were drawn from an equal number of edges selected uniformly at random from all possible edges in the connectome.
2. *Random FC-*Matched: Synergy values were drawn from edges selected uniformly at random but with the constrain that their original FC weights (or redundancy values) were comparable to those of the actual internal edges being analyzed. This controls for potential dependencies between synergy and FC/redundancy.
3. *Random Cycle Internal* : Synergy values were drawn after selecting a number of random cycles matching the number and lengths of original cycle. This test was performed to exclude the possibility that the origin of higher synergy is in the cyclic structure itself, rather than in the topological properties of the specific cycles in the connectome.

Statistical tests (two-sided t-tests) was used to compare the synergy distributions between border links, internal links, and the random distributions.

### N. Code Availability

All custom code and analysis scripts used for the scaffold computation, fingerprinting analysis, and information-theoretic calculations described in this manuscript are publicly available on GitHub at the following address: https://github.com/simonepoetto/topologicalfingerprinting.

## ACKNOWLEDGMENTS

G.P. acknowledges partial support by ERC Consolidator Grant RUNES (Grant no. 101171380) and the MSCA Doctoral Network *BeyondTheEdge*(Grant no. 101120085). A.S. acknowledges funding from the European Union’s Horizon 2020 research and innovation programme under the Marie Sklodowska-Curie grant agreement no. 101208090.

## Supplementary Information: The Topological Architecture of Brain Identity

### S.1. ROBUSTNESS OF IDENTIFIABILITY

To assess the identifiability of each subject across scanning sessions, we computed the Pearson correlation between the vectorized upper triangles of the connectivity matrices (or scaffolds) from Session 1 and all subjects’ matrices in Session 2, and vice versa. Identification was considered correct when a subject’s own scan yielded the highest similarity across sessions. We computed success rates in both directions—Session *→*1 Session *→*2 and Session 2 → Session 1—and report the final identifiability as the average of the two. This bidirectional approach provides a robust estimate of cross-session consistency in individual-specific brain signatures.

### S.1. Robustness across atlases

Across all parcellations, *homological scaffolds* consistently outperform their density-matched (1%) thresholded FC counterparts, yielding higher identification success rates and larger withinvs. between-subject effect sizes (Table S1, Fig. S1, Table S2). For the Schaefer 300 atlas (Table S1), scaffolds achieve perfect identification in both sessions, while full FC performance plateaus around 90%. These results confirm that scaffolds retain strong subject-specific features even in extremely sparse representations.

Performance across different parcellation schemes (Table S2) further supports the robustness of scaffold-based fingerprinting. While scaffolds consistently outperform 1% thresholded FC for all atlases, their absolute accuracy varies with parcellation resolution. For coarse atlases like AAL-89 and Schaefer-100, scaffold performance is relatively modest (43% and 64.5%, respectively), but this is likely due to the small number of regions limiting the number and diversity of topological cycles. As the resolution increases (e.g., Shen 268, Schaefer 300, Glasser 360), scaffold performance rapidly rises, reaching near-perfect identifiability (99.5–100%). This trend supports the idea that the discriminative power of scaffolds emerges from capturing mesoscale topological features, which become increasingly expressive at higher resolutions.

### S.3. Localization of identifying edges

To explore where the most discriminative edges are located, we also computed identifiability success rates when restricting the analysis to within-network edges only (Table S3). As expected from prior work, FC-based identifiability is highest within canonical networks such as the Default Mode (73%) and Somatomotor (62%) systems. In contrast, scaffold-based performance drops substantially when limited to intra-network connections (e.g., 33% in DMN, 50% in SMN), confirming that their fingerprinting power does not rely on localized connectivity. Rather, as discussed in the main text, scaffold identity signatures are distributed across inter-network edges and capture subject-specific integration patterns across the entire brain. This supports a fundamentally different mechanism for individual differentiation, one that emphasizes topological integration over local coherence.

**TABLE S1.** Identifiability (success rates) for the Schaefer 300 atlas with full time series.

|  | FC full | FC 25% | FC 1% | Scaffolds |
| --- | --- | --- | --- | --- |
| Session 1 | 89% | 90% | 87% | **100%** |
| Session 2 | 92% | 90% | 88% | **100%** |

**TABLE S2.**
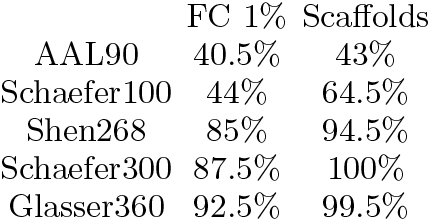
Identifiability (success rates) for the various atlases of 1% thresholded FC vs. scaffolds.

|  | FC 1% | Scaffolds |
| --- | --- | --- |
| AAL90 | 40.5% | 43% |
| Schaefer100 | 44% | 64.5% |
| Shen268 | 85% | 94.5% |
| Schaefer300 | 87.5% | 100% |
| Glasser360 | 92.5% | 99.5% |

**FIG. S1.**
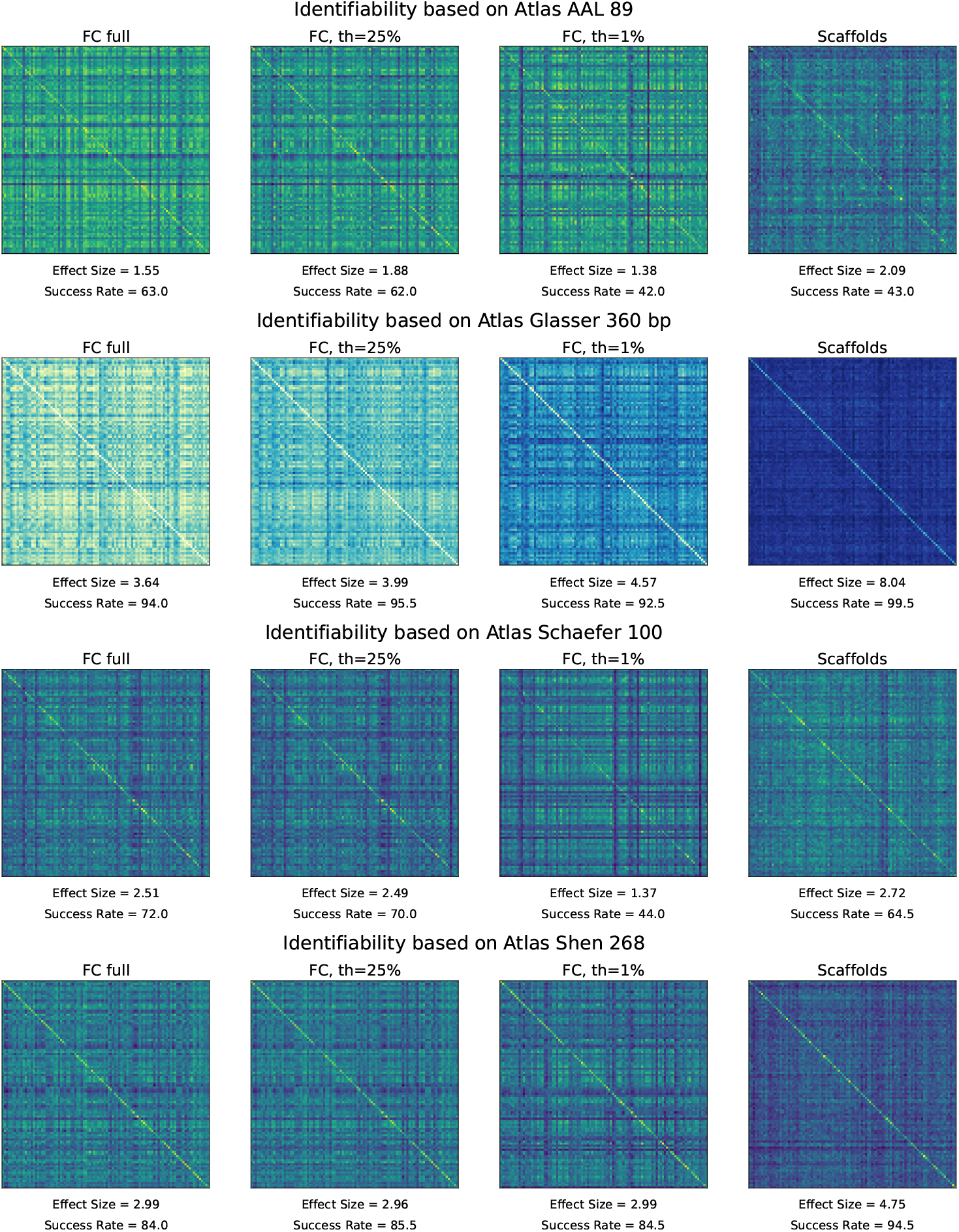
Similarity matrices for different atlases and different FC thresholds versus scaffolds. From top to bottom, atlas and corresponding number of regions: AAL (89), Glasser (360), Schaefer (100), Shen (268).

**TABLE S3.** Success rate when considering within-network connections only.

|  | FC | Scaffolds |
| --- | --- | --- |
| Frontoparietal | **40%** | 21% |
| Default | **73%** | 33% |
| DorsalAtt | **25%** | 10% |
| Limbic | **24%** | 10% |
| VentralAtt | **31%** | 19% |
| Somatomotor | **62%** | 50% |
| Visual | **50%** | 33% |

**FIG. S2.**
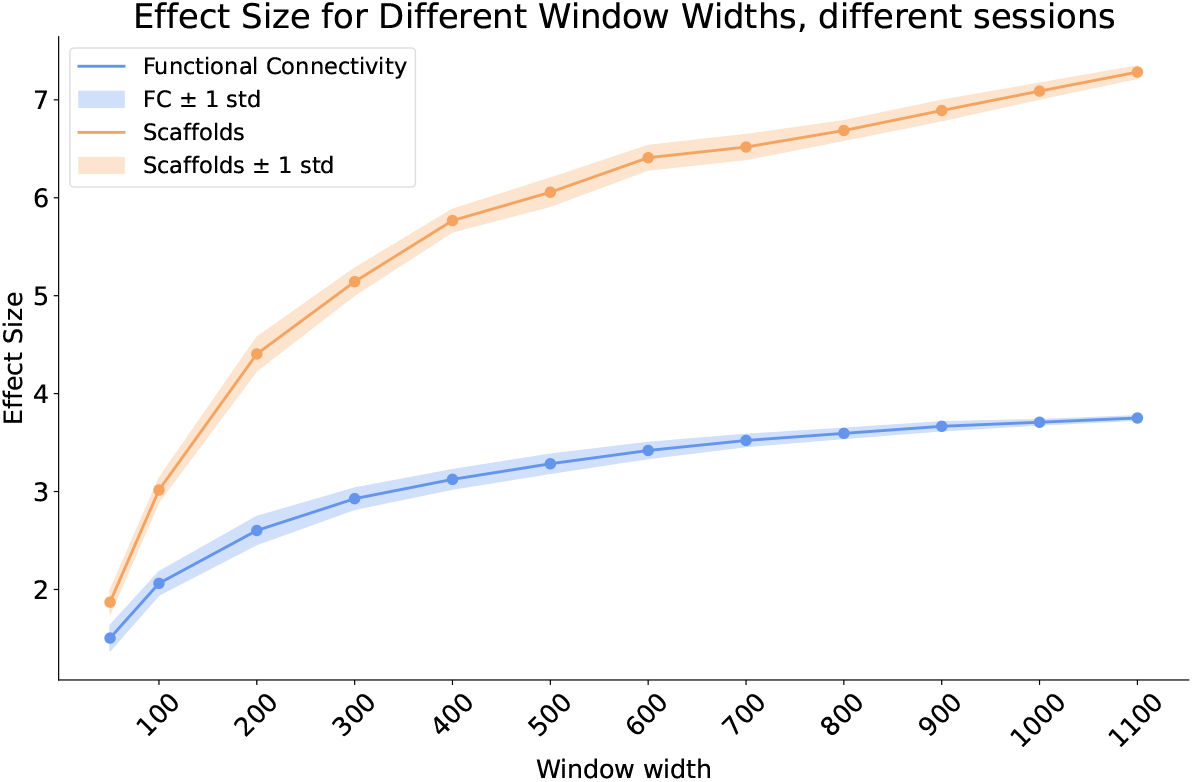
Effect sizes of intra-vs inter-subject similarities. We show here results for FC (1%) and scaffolds as a function of window width, complementary to Fig. 2D.

### S.2. EFFECTS OF PREPROCESSING

Table S4 reports the impact of several standard preprocessing steps on fingerprinting accuracy across different network representations. Homological scaffolds consistently outperform all functional connectivity (FC) benchmarks, including full, 25%, and 1% thresholded matrices. In most conditions, scaffolds achieve near-perfect identification (99–100%), demonstrating remarkable robustness to changes in preprocessing. Notably, scaffold performance remains high whether only LR phase encoding is used or LR/RL runs are averaged, and is largely unaffected by the inclusion or omission of global signal regression (GSR) or high-pass filtering. The only substantial performance drop is observed when low-pass filtering is omitted: in this setting, scaffold accuracy decreases to 83.5%, though it still exceeds all FC-based methods under the same condition. This suggests that low-pass filtering helps suppress high-frequency noise that may interfere with the topological cycle formation central to scaffold construction. By contrast, FC performance is more variable and systematically lower across all preprocessing variants. These findings highlight the superior stability of scaffold-based fingerprinting under typical sources of preprocessing variability, making it especially promising for robust subject identification across heterogeneous fMRI pipelines.

**TABLE S4.**
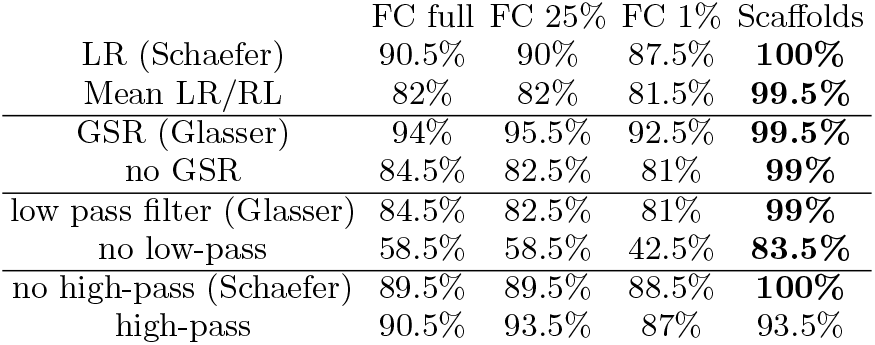
Effects of Preprocessing effects on identifiability (success rates).

|  | FC full | FC 25% | FC 1% | Scaffolds |
| --- | --- | --- | --- | --- |
| LR (Schaefer) | 90.5% | 90% | 87.5% | **100%** |
| Mean LR/RL | 82% | 82% | 81.5% | **99.5%** |
| GSR (Glasser) | 94% | 95.5% | 92.5% | **99.5%** |
| no GSR | 84.5% | 82.5% | 81% | **99%** |
| low pass filter (Glasser) | 84.5% | 82.5% | 81% | **99%** |
| no low-pass | 58.5% | 58.5% | 42.5% | **83.5%** |
| no high-pass (Schaefer) | 89.5% | 89.5% | 88.5% | **100%** |
| high-pass | 90.5% | 93.5% | 87% | 93.5% |

### S.3. TRAIT VS. STATE SENSITIVITY IN SCAFFOLD AND FC REPRESENTATIONS

In addition to cross-session identifiability, we examined fingerprinting performance within a single scanning session (i.e., intrasession comparisons). Results are reported in Figures S3. In this setting, both functional connectivity (FC) and scaffold-based representations show improved success rates and larger effect sizes relative to intersession analyses, as expected due to reduced temporal variability. However, we observe that –for short timescales– the gap between FC and scaffold performance narrows substantially: the effect sizes and success rates for FC increase and become more comparable to those of scaffolds (Fig. S4).

This convergence suggests an important interpretive distinction between the two representations. The scaffold, by design, captures the persistent topological structure of functional interactions—features that are stable over time and thus likely reflect individual traits in brain organization. FC, in contrast, appears more sensitive to transient fluctuations in co-activation patterns that may reflect cognitive or physiological states present during scanning. As a result, while scaffolds maintain high identifiability across sessions (trait-like stability), FC gains relative discriminability when constrained to a single session where state-related variability is minimized.

This interpretation aligns with the emerging view that functional fingerprinting reflects a mixture of stable, trait-like architecture and more dynamic, state-driven reconfigurations. Under this light, scaffolds may therefore provide a complementary lens on brain individuality, emphasizing temporally invariant mesoscale integration patterns less influenced by momentary fluctuations in cognitive or physiological state.

Finally, these results also align with reports that *instantaneous scaffolds* (built from higher-order timeseries rather than FC as in our case) [35] do not show high individual fingerpriting capacity, but rather excel at task decoding, pointing to different distributed structures captured by different scaffold constructions at different temporal scales.

**FIG. S3.**
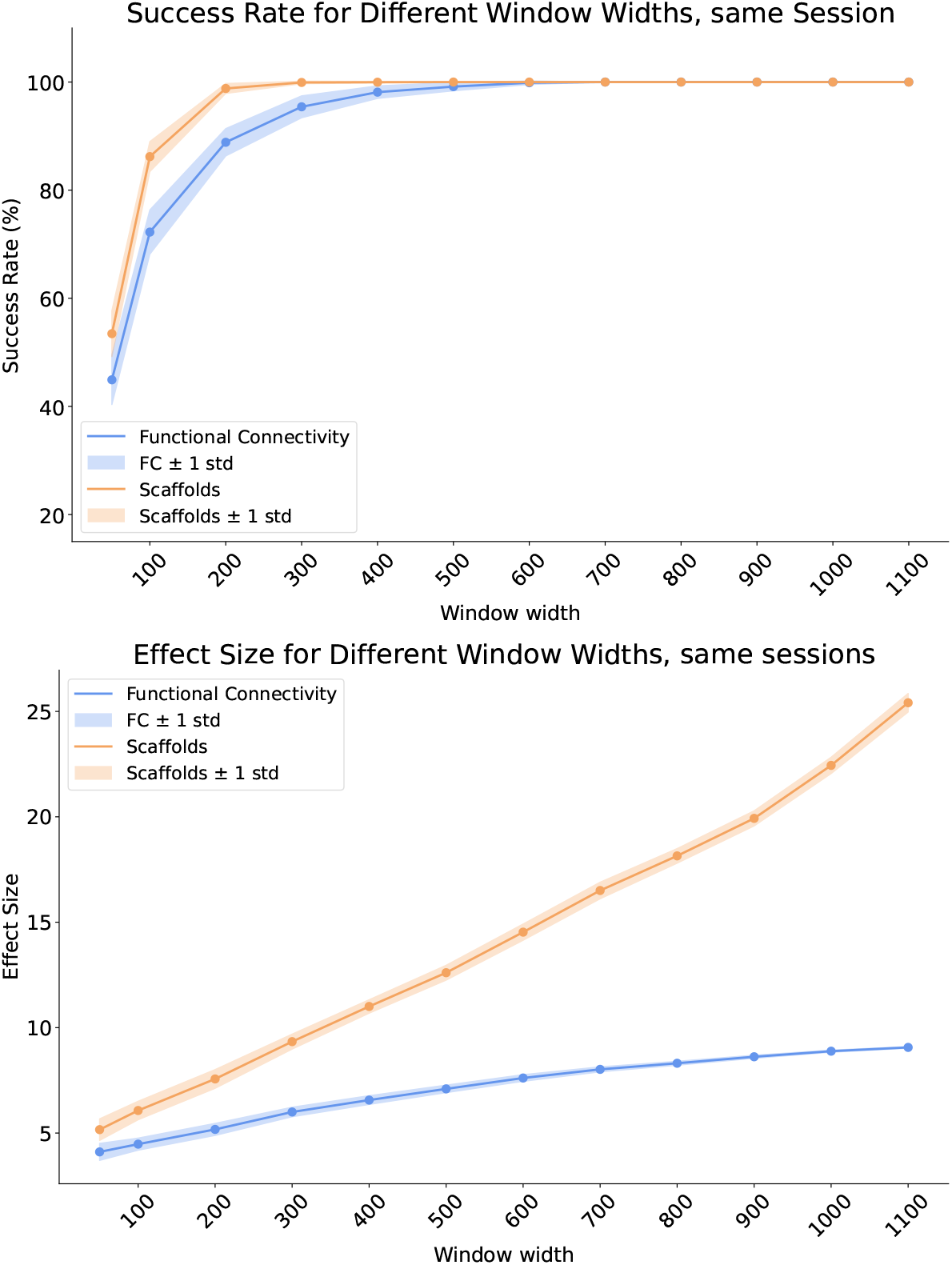
Intrasession fingerprinting performance. Comparison of identifiability success rates for FC and scaffold representations when computed within-session (left) and across-sessions (right).

**FIG. S4.**
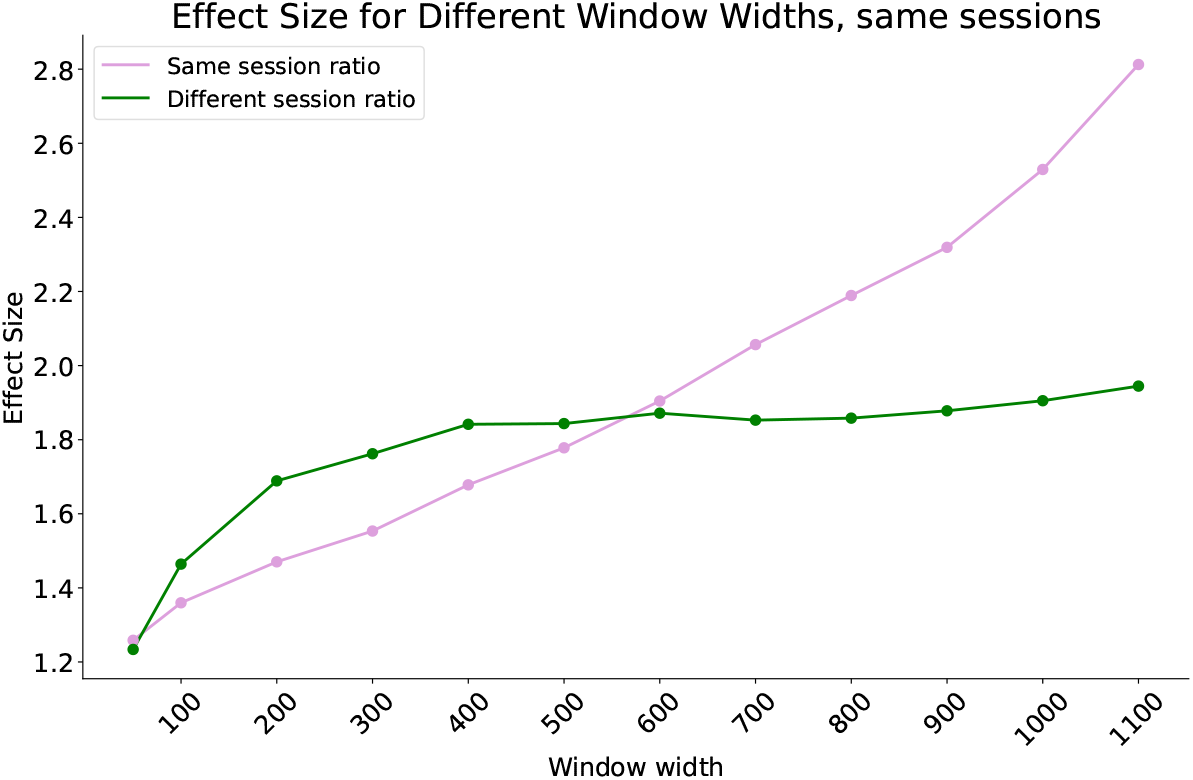
Ratio of scaffold vs FC effect sizes, intra- and inter-session as a function of the window length. We see that for short time scales within the same sessions the ratio between effect sizes for scaffolds and FC is quite small, signaling a comparable capacity of FC and scaffolds to fingerprint.

